# Generation of functional adipoids from human induced pluripotent stem cells

**DOI:** 10.64898/2026.10.30.762496

**Authors:** Laura C. Hinte, Ryoko Okamoto, Sophie Jansen, Stefan E. Spirig, Sofia Tsiepa, Mitsugu Shimobayashi, Michael N. Hall

**Affiliations:** Institute of Human Biology (IHB), Pharma Research and Early Development (pRED), F. Hoffmann-La Roche Ltd, Basel, Switzerland; 360 Labs Genomics Platform, Pharma Research and Early Development (pRED), F. Hoffmann-La Roche Ltd, Basel, Switzerland; Biozentrum, University of Basel Basel, Switzerland

## Abstract

Human adipose tissue models are essential for metabolic research. Traditional primary cell cultures are poorly scalable and not suitable for drug screening or modeling of genetic disease. Conventional two-dimensional adipocyte cultures cannot withstand the long-term treatment required to mimic chronic conditions such as obesity and insulin resistance. To overcome these limitations, we developed an accessible protocol to produce three-dimensional human induced pluripotent stem cell (hiPSC)-derived adipocyte microtissues, termed adipoids. hiPSC-derived adipoids display long-term stability and viability. Single-nucleus RNA sequencing revealed that adipoids contain mature adipocytes and adipocyte stem/progenitor cells, similar to human adipose tissue. Furthermore, hiPSC-derived adipoids contain unilocular adipocytes, exhibit stimulated lipolysis, and secrete adipokines. Thus, hiPSC-derived adipoids provide a functional, scalable and genetically tractable model to perform large-scale pharmacological screens and to investigate human metabolic disease.

## Introduction

Perturbed white adipocyte function, such as alterations in the capacity to store and release lipids, is associated with cardiometabolic disease^1,2^. To gain mechanistic insights into adipocyte (dys-)function, *in vitro* systems with mature adipocytes are required. Unfortunately, the buoyancy and fragility of mature adipocytes in conventional two-dimensional (2D) culture impede mechanistic studies^3^. The development of reliable *in vitro* models of human adipose tissue is essential for understanding adipocyte function and deciphering the underlying mechanisms of metabolic diseases^4,5^.

Historically, such *in vitro* models have predominantly relied on immortalized cell lines or primary adipocyte stem/progenitor cells (ASPCs) isolated from the stromal vascular fraction (SVF) of adipose tissue^6^. While these systems have provided foundational insight, they present significant limitations. Primary ASPCs are often restricted in availability and suffer from limited replicative capacity, posing scalability challenges for long-term metabolic studies. Regarding immortalized cell lines, most accessible lines have been generated from patients with genetic diseases^7,8^, thereby introducing confounding factors. Furthermore, the established models derived from mesenchymal stem cells are not well-suited for efficient genetic manipulation^9^, hindering the investigation of specific disease-associated mutations.

To circumvent these limitations, human induced pluripotent stem cells (hiPSCs) have emerged as a versatile alternative. hiPSCs have been successfully differentiated into adipocytes in 2D monolayer cultures^10,11^, but these 2D adipocyte cultures are difficult to maintain over extended periods. As adipocytes mature and accumulate lipid droplets, they frequently detach from the culture surface, abruptly terminating experiments^3^. This fragility and buoyancy are problematic for modeling chronic metabolic dysfunctions, such as obesity and insulin resistance. Several groups have published protocols for three-dimensional (3D) adipocyte culture^12–16^. However, these 3D culture protocols rely mostly on primary ASPCs, constraining scalability and genetic tractability.

To address these current challenges, we developed an easy-to-execute, highly reproducible protocol to generate 3D adipocyte microtissues, termed adipoids, from human hiPSCs. We characterized the biological functionality and cellular composition of these adipoids using single-nucleus RNA sequencing (snRNAseq) and benchmarked them against primary ASPC-derived adipoids. Our hiPSC-derived adipoid model offers an accessible system for large-scale pharmacological screens and the investigation of chronic metabolic diseases and genetic lipid disorders.

## Results

### Generation of adipoids from hiPSCs

To establish a scalable adipose model, we developed a directed differentiation protocol to produce adipoids, which we applied to three distinct hiPSC lines, namely KOLF2.1J (KOLF)^17^, SCTi003-A (SCT) from STEMCELL Technologies and WTC-11 (WTC)^18^. We first generated mesenchymal progenitor cells (MPCs) using a defined differentiation medium (**Fig. 1a** and **1b**). After 21 days of culture and expansion, cells displayed a characteristic spindle-like morphology (**Fig. 1c**). These MPCs could be expanded further or cryopreserved. For generation of adipoids, the MPCs were seeded on ultra-low attachment plates to form MPC spheroids which were subsequently embedded in a basement membrane matrix (Geltrex). Consistent with previous observations with primary ASPCs^14,16^, MPC spheroids exhibited characteristic cellular protrusions within days of embedding. MPCs were then subjected to 18 days of adipogenic differentiation to generate adipoids (see Materials and Methods). Adipoids were maintained in serum-free medium supplemented with low glucose (5 mM) and low insulin (10 nM) with half medium changes every 3-4 days. Adipoids from all three hiPSC lines could be maintained for at least 90 days in culture without noticeable signs of cell degeneration or death. Adipoids consistently contained mature, primarily unilocular adipocytes approximately 50 µm in diameter by day 60 (**Fig. 1d**), indicating the robustness and applicability of the protocol.

**Figure 1.**
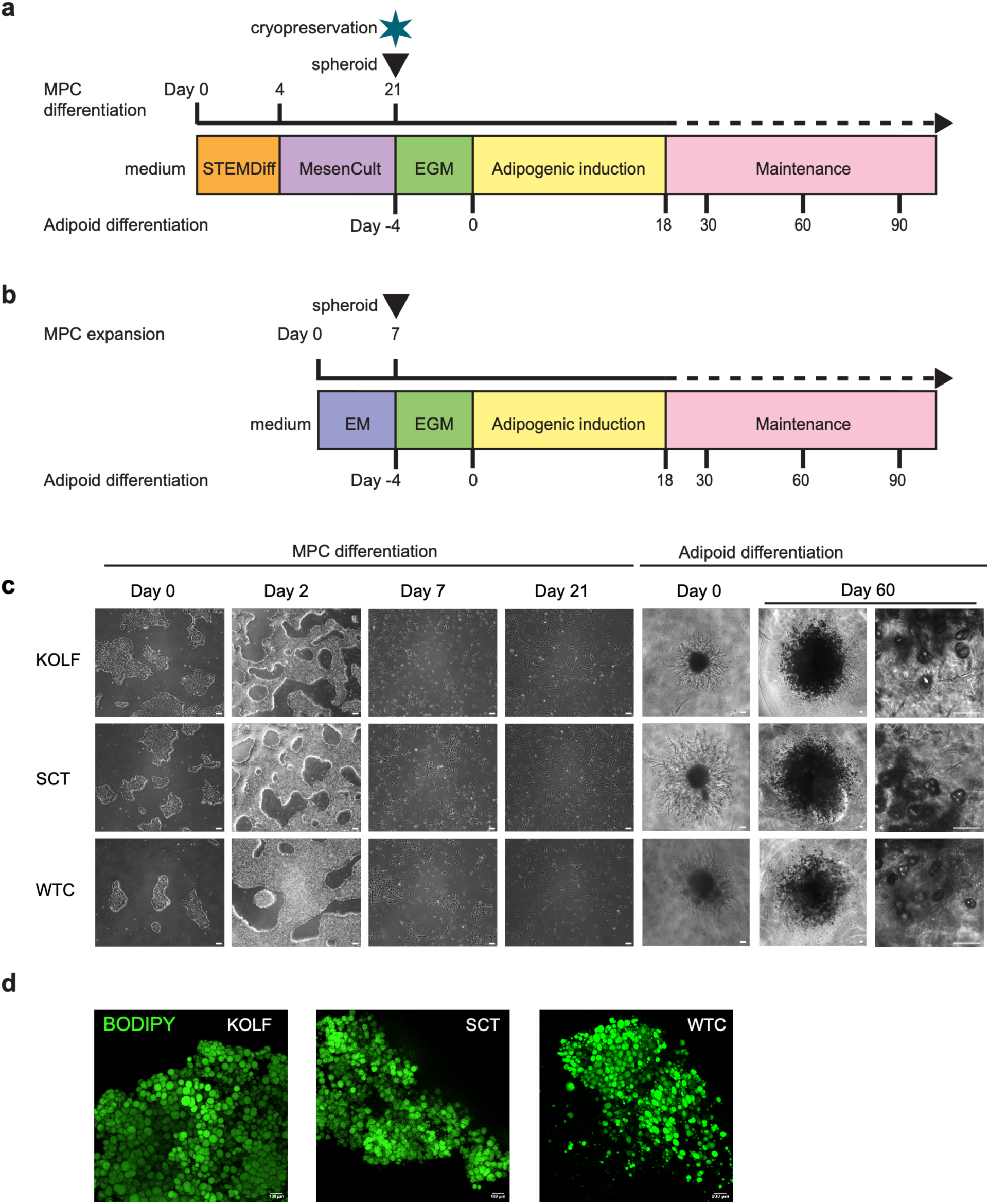
Generation of adipoids from hiPSCs. **a**, Generation of MPCs from hiPSCs and subsequent generation of adipoids. STEMDiff, STEMdiff-ACF Mesenchymal Induction Medium. MesenCult, MesenCult-ACF Medium. EGM, Endothelial Cell Growth Medium-2. **b**, Generation of adipoids from cryopreserved hiPSC-derived MPCs. EM, Expansion Medium. **c**, Representative microscopic images from indicated days of MPC and subsequent adipoid generation across three hiPSC lines. Images at day 60 were taken of cleared adipoids. Scale bar is 100 µm. **D**, Representative (partial) images of mature adipoids stained with BODIPY (green). Scale bar is 100 µm.

### hiPSC-derived adipoids contain stromal cells and mature adipocytes

We next characterized the cellular composition of the hiPSC-derived adipoids. We isolated nuclei from frozen adipoids at day 60 post adipogenic differentiation and subjected them to single nucleus RNA sequencing (snRNAseq) (**Fig. 2a**). 4,694 nuclei in total were sequenced from the three hiPSC lines (**Supplemental Fig. S1a**). Adipoids from all three hiPSC lines contained two major cell types (**Fig. 2b**) which we could annotate as adipocytes and ASPCs, based on expression of canonical markers (**Fig. 2c-2f).** Interestingly, cells identified as ASPCs expressed adipocyte markers such as *GPAM* and *ADIPOQ* but without *PLIN1* encoding the major structural protein for lipid droplets Perilipin1, suggesting that these cells are differentiating toward adipocytes. We detected little-to-no expression of pluripotency markers (*KLF4*, *NANOG*, *POU5F1*, and *SOX2*) (**Supplemental Fig. S1b**), indicating that the adipoids were free of residual hiPSCs. Furthermore, the cellular landscape was similar across all three hiPSC lines (**Fig. 2g**). These data demonstrate the reliability of our directed differentiation protocol in producing adipoids from hiPSCs.

**Figure 2.**
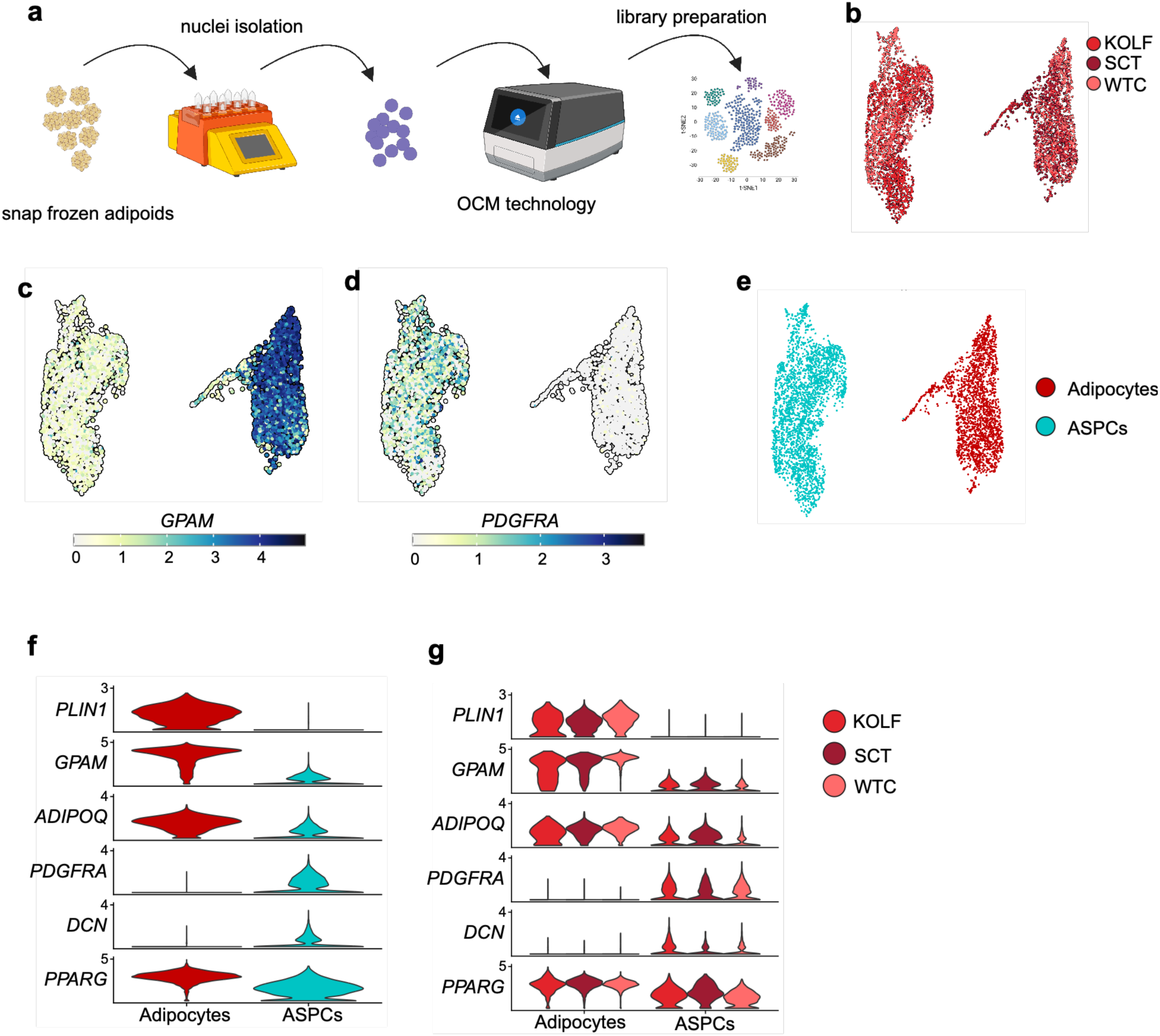
Characterization of hiPSC-derived adipoids by snRNAseq. **a**, Workflow of single-nucleus RNA sequencing from adipoids. Nuclei are isolated from snap frozen adipoids and loaded onto 10x Genomics chips using on chip multiplexing (OCM) technology. Thereafter, libraries are prepared, sequenced and analyzed. **b,** Uniform manifold approximation and projection (UMAP) of 4,694 nuclei colored by hiPSC line. **c**, Normalized Expression of *GPAM,* a marker gene for adipocytes. **d**, Normalized Expression of *PDGFRA,* a marker gene of ASPCs. **e**, UMAP of cell type assignment. **f**, Gene expression of adipocyte and ASPC marker genes across assigned cell types. **g**, Gene expression of adipocyte and ASPC marker genes across assigned cell types and hiPSC lines.

### hiPSC-derived adipoids are transcriptionally similar to primary ASPC-derived adipoids

To benchmark the hiPSC-derived adipoids, we compared snRNAseq profiles of the hiPSC-derived adipoids and primary ASPC-derived adipoids^12–16^. To this end, we first expanded ASPCs from stromal vascular fraction (SVF) obtained from abdominal subcutaneous adipose tissue biopsies procured from abdominoplasty on three donors (**Table S1**). The expanded ASPCs were used to produce adipoids from which we isolated nuclei for snRNAseq analyses (see Materials & Methods). snRNAseq analyses revealed that hiPSC- and primary ASPC-derived adipoids display similar cellular composition consisting of mature adipocytes and ASPCs (**Fig. 3a** and **3b**). Having observed similar cellular composition of hiPSC- and primary ASPC-derived adipoids, we analyzed overall transcriptional similarity. We aggregated adipoid snRNAseq counts by cell type (adipocyte or ASPC) and source (hiPSC or donor) from which we calculated a Pearson’s correlation coefficient (R). We note that sex chromosome-linked genes were excluded from the analysis to avoid sex-related variability. Transcriptomic profiles of adipocytes and ASPCs were highly similar among adipoids derived from the three different hiPSC lines (R = 0.92-0.96 for adipocytes and 0.90-0.96 for ASPCs) (**Fig. 3c-d**), confirming the reliability of our protocol in producing adipoids from different hiPSC lines. More importantly, adipocytes and ASPCs in hiPSC- and primary ASPC-derived adipoids showed high similarity (R = 0.85-0.94 for adipocytes and 0.85-0.89 for ASPCs) (**Fig. 3c-d**). Thus, our directed differentiation protocol produces hiPSC-derived adipoids that faithfully reproduce primary ASPC-derived adipoids.

**Figure 3.**
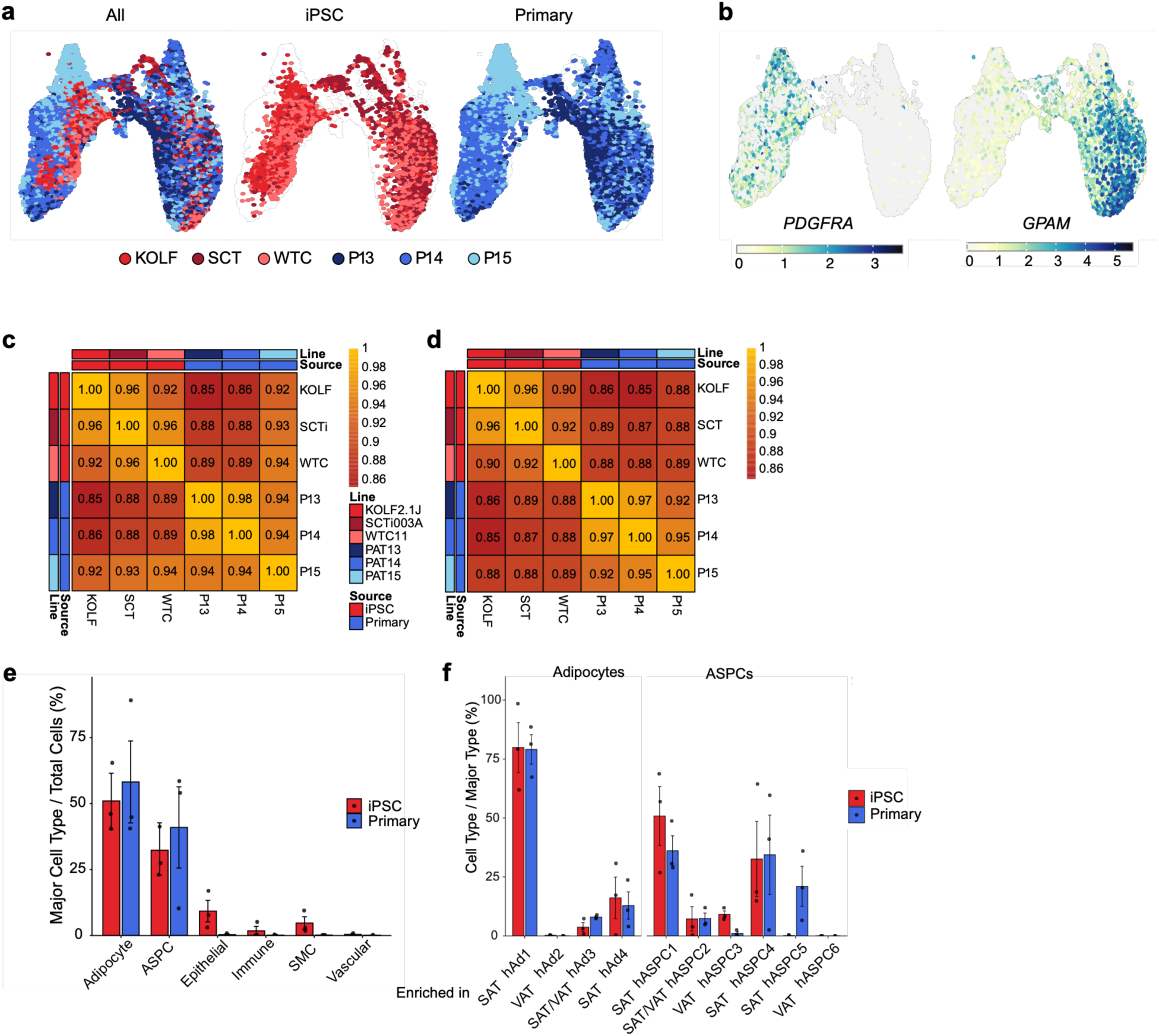
Transcriptional comparison of hiPSC- and primary ASPC-derived adipoids against human adipose tissue atlas. **a**, UMAP visualization of 17,577 nuclei colored by hiPSC lines or ASPC donors. **b**, Expression of *PDGFRA* and *GPAM*. **c**, Pearson’s R of adipocyte transcriptome among adipoids generated from hiPSCs or ASPCs isolated from tissue donors (P13, P14, P15). **d**, Pearson’s R of ASPC transcriptome among adipoids as in c. **e**, Proportion of cell types assigned by reference mapping against Emont *et al.* ^19^ in adipoids. **f**, Proportion of adipocyte and ASPC subclusters defined by Emont *et al.* ^19^ in adipoids.

### hiPSC-derived adipoids contain adipocytes and ASPCs resembling human subcutaneous adipose tissue

We next asked whether the subtle differences in gene expression in different adipoids reflect differences in cellular subpopulations. To address this question, we reference-mapped our adipoid snRNAseq datasets against published white human adipose tissue snRNAseq data^19^ and determined mapping and maximum prediction scores (**Supplemental Fig. S2a-2c**). The mapping score is a measure of how well individual cells in adipoids correspond to cells in human adipose tissue, independent of cell type. The maximum prediction score measures the confidence of the specific cell type assigned to a cell in adipoids. Adipocytes and ASPCs in hiPSC-derived and primary ASPC-derived adipoids were assigned to the same cell types in human adipose tissue based on high mapping and prediction scores (**Supplemental Fig. S2a-2c**). Of note, hiPSC-derived adipoids exhibited higher mapping scores than primary ASPC-derived adipoids (**Supplemental Fig. S2a**), suggesting that hiPSC-derived adipoids contain adipocytes and ASPCs more similar to the same cell types in human adipose tissue compared to primary ASPC-derived adipoids.

A previous study demonstrated that human white adipose tissue contains seven adipocyte (hAd1-7) and six ASPC (hASPC1-6) subpopulations^19^. To determine the nature of the adipocytes and ASPCs in adipoids, we mapped our snRNAseq data to the subpopulations of adipocytes and ASPCs in the published reference sets^19^. Overall, hiPSC- and primary ASPC-derived adipoids contain similar distribution of adipocyte and ASPC subpopulations (**Fig. 3e**). hiPSC-derived adipoids contain hAd1, hAd3, and hAd4 subpopulations that are enriched mainly in subcutaneous adipose tissue (SAT)^19^ and lack subpopulations such as hAd2 which are mainly found in visceral adipose tissue (VAT)^19^. Likewise, hiPSC-derived adipoids contain SAT-enriched ASPC subpopulations (hASPC1, hASPC2, hASPC4) and have little-to-no VAT-enriched subpopulation hASPC3 and 6 (**Fig. 3f**). Interestingly, hiPSC-derived adipoids lack the hASPC5 subpopulation, which was reported to be increased in individuals with high BMI (>40), compared to individuals with low BMI (<30)^19^. These findings suggest that hiPSC-derived models represent "healthy" SAT, possibly due to the adipoids having been maintained in physiological low glucose and low insulin conditions. Taken together, we conclude that our protocol produces hiPSC-derived adipoids that phenocopy the cellular heterogeneity and transcriptional profiles of adipocyte and ASPC subpopulations in human SAT (**Fig. 3f**).

The above reference mapping revealed that both hiPSC- and primary ASPC-derived adipoids consisted of approximately 50% adipocytes and 50% other type cells (**Fig. 3e**). A subset of cells in hiPSC-derived adipoids mapped to neither adipocyte or ASPC identities (**Fig. 3e**). This suggests that a small fraction of non-adipocyte cells may retain multi-lineage potential, possibly toward chondrogenic or myogenic lineages. Consistent with this possibility, *ACAN*, a marker gene typically associated with chondrogenic or myogenic lineages, was expressed in a subset of ASPCs in hiPSC-derived adipoids (**Supplemental Fig. S2d**).

### Adipoids contain functional adipocytes

Having established that our adipoids are transcriptionally similar to primary ASPC-derived adipoids, we next characterized the functionality of hiPSC-derived adipoids. The primary function of adipocytes is to store excess energy and to release it upon beta-adrenergic stimulation^1^. Histological analysis revealed that the hiPSC-derived adipoids accumulate unilocular lipid droplets similar to adipoids produced from primary ASPCs (**Fig. 4a**). Beta-adrenergic stimulation with isoproterenol increased glycerol release in both hiPSC- and primary ASPC-derived adipoids (**Fig. 4b**), suggesting intact energy storage and release functions of adipocytes. Furthermore, hiPSC-derived adipoids, like primary ASPC-derived adipoids, secreted the well-established adipokines adiponectin and leptin (**Fig. 4c** and **4d**), supporting the maturity of the hiPSC-derived adipoids. In summary, our directed differentiation protocol produces functional adipoids.

**Figure 4.**
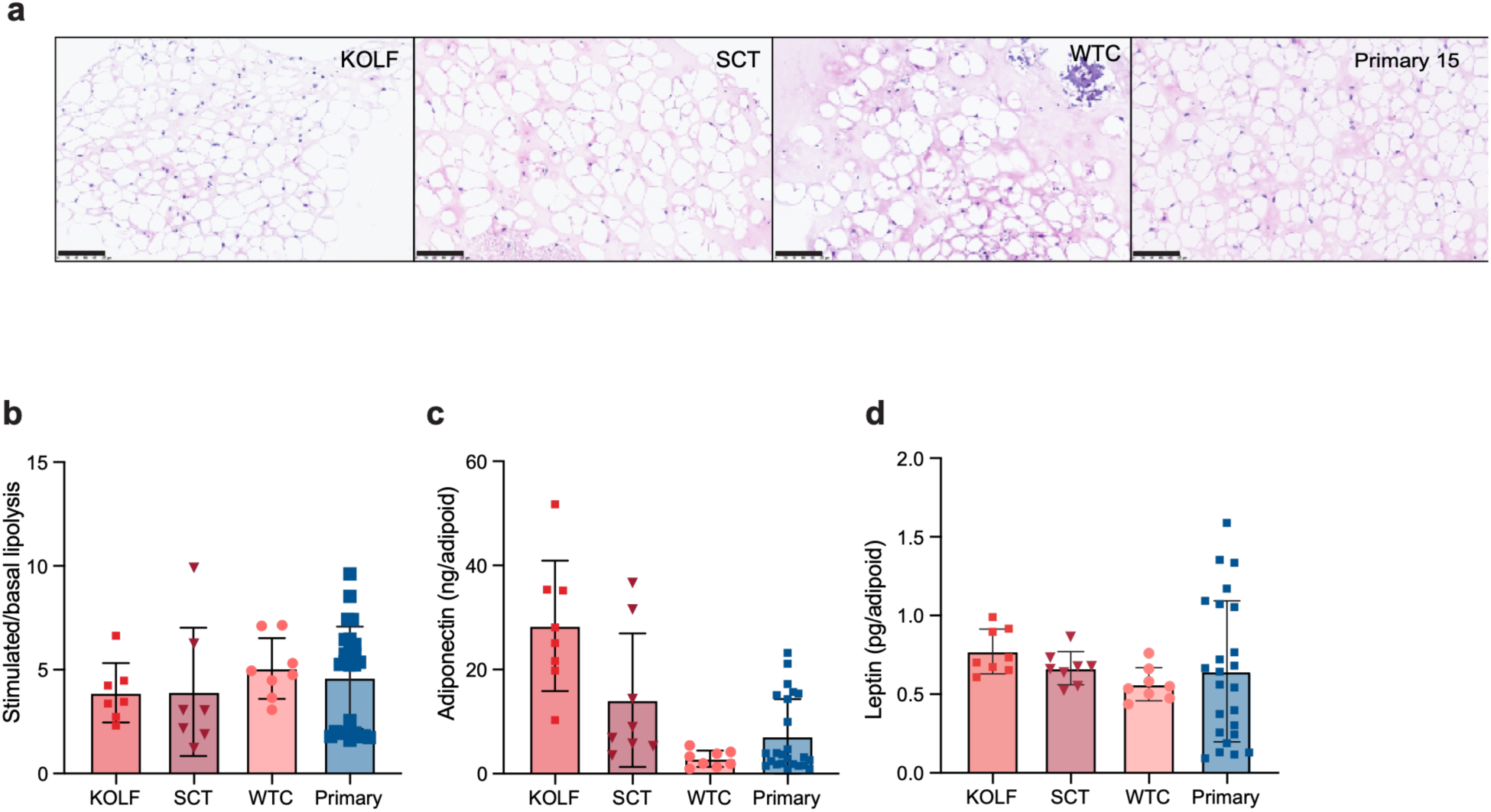
Functional characterization of hiPSC-derived adipoids. **a**, Representative images of histological sections stained with hematoxylin and eosin of adipoids from hiPSCs and one primary donor. Scale bar is 100 µm. **b**, Stimulated over basal lipolysis. One dot represents one adipoid (n= 8 for hiPSCs). For primary cell derived adipoids, data from three donors is presented. One dot represents one adipoid. **c**, Adiponectin measured in cell culture supernatant. Supernatant from three adipoids was pooled per replicate. **d**, Leptin measured in cell culture supernatant. Supernatant from three adipoids was pooled per replicate.

## Discussion

Deregulated function of adipose tissue is central to the etiology of metabolic disease in humans. However, progress in understanding underlying mechanisms has been impeded by a lack of suitable, genetically tractable, and stable *in vitro* human adipose tissue models. In this study, we have developed and characterized a robust hiPSC-derived 3D adipoid system. Our hiPSC-derived adipoids exhibit robust functional maturity - marked by stimulus-responsive lipolysis and adipokine secretion - and display cellular heterogeneity similar to primary ASPC-derived adipoids and human adipose tissue.

There are several key steps in our protocol to produce functional and stable adipoids from hiPSCs. First, embedding MPC spheroids in a basement membrane matrix (Geltrex) was strictly required. In the absence of a matrix, MPC spheroids and differentiated adipoids failed to maintain structural integrity. Second, embedding prevented lipid-induced buoyancy (detachment) observed in conventional 2D systems, allowing long-term continuous culture (at least 90 days). Lastly, after adipogenic differentiation, physiological conditions (5 mM glucose and 10 nM insulin) maintained functionality of adipoids, again allowing long-term culture.

Long-term continuous culture conditions offer a possibility to model chronic metabolic disease. For example, it is of interest to examine whether long-term culture of hiPSC-derived adipoids in pathological conditions – e.g. high insulin, high glucose, and/or high lipid conditions – cause phenotypes similar to metabolic disease such as excessive lipid accumulation and insulin resistance.

Another advantage of hiPSC-derived adipoids over conventional primary cells is their genetic tractability. Unlike primary cells, hiPSCs allow genome engineering such as CRISPR/Cas9-mediated editing. This in turn allows seamless introduction of patient-specific mutations, providing an unprecedented tool for modeling human genetic diseases such as lipodystrophies^20^. Furthermore, the inherent scalability of hiPSCs allows high-throughput pharmacological screening.

While our model overcomes significant hurdles in accessibility and scalability, some limitations remain. Like all *in vitro* systems, our adipoids lack the full complexity of the native tissue environment, including rhythmic nutrient supply and heterogeneous tissue environments, e.g. lack of vasculature, immune cells, and innervation.

Furthermore, snRNAseq analyses suggested that some cells in hiPSC-derived adipoids may retain the capacity to give rise to other lineages which are distinct from the native adipose-specific progenitors. Despite these limitations, hiPSC-derived adipoids can be a valuable addition to current cell systems in the field of adipocyte research and will serve as a basis to fully model the complex human adipose tissue.

## Materials and methods

### Ethics declaration

This study used the previously derived hiPSC lines KOLF2.1J (RRID:CVCL_B5P3), SCTi003-A (STEMCELL Technologies, #200-0511, RRID:CVCL_C1W7), and WTC-11 (RRID:CVCL_Y803) . All lines were derived under local ethical approval at the original derivation sites and with informed consent from the donors. No new hiPSC lines were derived in this study. The Institute of Human Biology (Roche) and the Hall laboratory hold the ethical approval required to work with hiPSC lines. SVF cells were isolated from human abdominal subcutaneous adipose tissue obtained from cosmetic surgery. Tissues and associated clinical information were obtained from patients undergoing elective aesthetic surgery who had no diabetes or other known disease. All tissues were sourced under contractual agreements that comply with applicable legal and ethical regulations. The study was conducted in accordance with the Declaration of Helsinki and was reviewed and approved by the Direction Générale de la Recherche et de l’Innovation of the Ministère de l’Enseignement Supérieur et de la Recherche (France), Ethics Committee No. AC-2023-6079.

### Cell culture

All cells were cultured at 37 °C, 5% CO2 and 90% relative humidity.

### Isolation of the stromal vascular fraction

Adipose tissue pieces (1 cm × 1 cm × 1 cm) were soaked for 15 min at room temperature in 1% antibiotic-antimycotic (100×; Gibco, #15240062) in DPBS (Gibco, #14190144), washed in DPBS and transferred to a 50 ml tube. The tissue was minced with scissors in 16 ml pre-warmed digestion buffer (HBSS++ (Hanks’ balanced salt solution with Ca2+ and Mg2+; Gibco, #14025134) supplemented with 1% (w/v) BSA (Sigma-Aldrich, #A9418-100G)) until the total volume reached 26 ml. Then, 4 ml digestion buffer containing 15,000 units collagenase from Clostridium histolyticum (Sigma-Aldrich, #6885-5G) and 1,200 units DNase I (Merck, #34536282001) were added, and the tissue was digested at 37 °C for 40 min with shaking. The digest was resuspended by pipetting after 20 and 30 min to ensure complete digestion. Digestion was stopped by adding 400 µl of 0.5 M EDTA (Millipore, #324504) and pipetting up and down. Digests were centrifuged at 300 × g for 5 min. Floating adipocytes and lipids were collected with cut Pasteur pipettes, the supernatant was decanted, and the SVF pellet was resuspended in digestion buffer without enzymes. The SVF was filtered through a 100 µm cell strainer and centrifuged at 450 × g for 5 min. The pellet was resuspended in 5 ml ACK lysis buffer (Gibco, #A1049201) and incubated for 5 min at room temperature, and lysis was stopped by adding a 20 ml digestion buffer. Cells were centrifuged at 450 × g for 5 min, resuspended in expansion medium (DMEM/F-12 (Gibco, #31331028), 10% FBS (Gibco, #A5670402), 1% penicillin-streptomycin (Gibco, #15070063)) and counted. Cells were either plated for expansion in T75 flasks (Corning, #353136) or cryopreserved in CellBanker 1 (Nippon Zenyaku Kogyo, #11910). Freshly isolated or freshly thawed SVF cells (passage 0) were plated in expansion medium in T75 flasks. At 80% confluency, cells were passaged 1:5 into T150 flasks (Corning, #353112) using Animal Component-Free Cell Dissociation Kit TrypLE Express (Gibco, #12604-013). Cells from passage 2 were used to generate adipoids.

### Generation of MPCs from hiPSCs

All media were supplemented with 1% penicillin-streptomycin (Gibco, #15140-122). hiPSCs were thawed into wells of a Geltrex-coated (Gibco, #A14132-02) 6-well plate in mTeSR Plus medium (STEMCELL Technologies, #100-0276) supplemented with 10 µM ROCK inhibitor (Y-27632; STEMCELL Technologies, #72305), which was removed the next day. At 70-80% confluency, hiPSCs were passaged with 0.5 mM EDTA and seeded at 480,000 cells per well of a Geltrex-coated 6-well plate in mTeSR Plus with ROCK inhibitor (day −2). The ROCK inhibitor was removed the next day (day −1). On day 0, the medium was changed to STEMdiff-ACF Mesenchymal Induction Medium (STEMCELL Technologies, #05241) and was replaced daily. On day 4, the medium was changed to complete MesenCult-ACF Medium (STEMCELL Technologies, #05445 supplemented with #05446) and replaced daily until day 6. On day 6, cells were harvested with Gentle Cell Dissociation Reagent (STEMCELL Technologies, #100-0485), washed in complete MesenCult-ACF Medium, centrifuged at 300 × g for 8 min and plated at 100,000 cells per well of a collagen I-coated 6-well plate (Greiner Bio-One, #657950) to expand the developing MPCs. Half of the medium was changed daily until cells reached 80% confluency, usually at day 9. Cells were then harvested with the Animal Component-Free Cell Dissociation Kit (STEMCELL Technologies, #05426), and 300,000 cells were seeded into a collagen I-coated 175 cm2 flask (Corning, #356487). The medium was changed every 2-3 days, and cells were passaged at 80% confluency (usually day 17). On day 21, MPCs were cryopreserved in CellBanker 1 or used to generate adipoids.

Cryopreserved MPCs (300,000 cells) were thawed into expansion medium (EM) supplemented with 2.5 ng/ml FGF2 (PeproTech, #100-18B-1MG) in a collagen I-coated 175 cm2 flask and expanded for 1-2 passages before adipoid generation. Cells were passaged with TrypLE Express.

### Generation of adipoids

The adipoid protocol was developed based on published work^12,14,15^. Adipoids were generated from expanded SVF cells (passage 2) or expanded hiPSC-derived MPCs (passage 3 or 4). Cells were harvested with TrypLE Express, washed in Endothelial Cell Growth Medium-2 (EGM-2, Lonza, #CC-3162) and counted. Then, 20,000 cells were seeded per well of a 96-well ultra-low attachment plate (Corning, #7007), and the plates were centrifuged at 100 × g for 1 min and incubated at 37 °C for 2 days. The medium was then removed, and the spheroids were embedded in 30 µl Geltrex, incubated for 30 min at 37 °C, and overlaid with 200 µl EGM-2 medium. After 2 days, adipogenic induction was started (day 0) with a half medium change using induction medium I (Williams E Medium (Gibco, #12551032), 1× insulin-transferrin-selenium (Gibco, #41400045), 1× penicillin-streptomycin, 1× GlutaMAX (Gibco, #35050061), 100 nM dexamethasone (STEMCELL Technologies, #72092), 500 µM isobutylmethylxanthine (STEMCELL Technologies, #72762), 10 µM rosiglitazone (STEMCELL Technologies, #72624), 10 nM hydrocortisone (Sigma-Aldrich, #H6909), 2 nM 3,3′,5-triiodo-L-thyronine (STEMCELL Technologies, #100-0548), 33 µM biotin (MP Biomedicals, #02101023-CF) and 17 µM pantothenic acid (Sigma-Aldrich, #P5155)). Half of the medium was changed every 3-4 days, alternating between induction medium I and induction medium II (Williams E Medium, 1× insulin-transferrin-selenium, 1× penicillin-streptomycin, 1× GlutaMAX and 100 nM dexamethasone), for three cycles (18 days). On day 18, the medium was changed to maintenance medium (1:1 mix of low-glucose DMEM (Gibco, #11885084) and Ham’s F-12K (Kaighn’s) Medium (Gibco, #21127022), 1× GlutaMAX (Gibco, #35050061), 15 mM HEPES (Gibco, #15630056) and 1× penicillin-streptomycin) supplemented with 10 nM insulin (Sigma-Aldrich, #I9278-5ML). Half of the maintenance medium was changed every 3-4 days. For all experiments in this study, adipoids were maintained until day 60 after adipogenic induction.

### Histology and staining

Adipoids were washed once in DPBS on ice, fixed in 4% paraformaldehyde (Thermo Fisher Scientific, #J19943.K2) for 30 min at room temperature, washed twice in DPBS and stored at 4 °C.

### Hematoxylin and eosin staining

PFA-fixed adipoids were mounted in custom-made molds filled with pre-liquefied HistoGel (Epredia, #HG-4000-012). The molds were carefully lifted into biopsy cassettes, dehydrated overnight and embedded in paraffin using a HistoCore PEARL tissue processor (Leica). Blocks were sectioned at 3.5 µm on a Microm HM355S microtome (Thermo Fisher Scientific) and transferred onto TOMO IHC adhesive glass slides (Roche, #07098928001). Slides were dried overnight at 40 °C. Hematoxylin and eosin staining was performed on a Ventana HE 600 (Roche), and sections were imaged with a NanoZoomer S360 (Hamamatsu).

### Confocal imaging

Formalin-fixed adipoids were stained with 2 µM BODIPY 493/503 (Thermo Fisher Scientific, #D3922) and 1 µg/ml DAPI (Thermo Fisher Scientific, #62248) in DPBS for 48 h at 4 °C, washed three times in DPBS and transferred to an optical-bottom microplate (Thermo Fisher Scientific, #165305). Thirty minutes before imaging, 20-30 µl clearing solution (60% (v/v) glycerol (Sigma-Aldrich, #G5516-100ML) and 2.5 M fructose (Sigma-Aldrich, #F0127-10MG)) was added to each adipoid. Adipoids were imaged on a CSU-W1 spinning disk microscope (Nikon).

### Lipolysis assay

Adipoids were washed once in DPBS, transferred to a new plate and starved of insulin and glucose overnight in assay medium (phenol red-free, glucose-free DMEM (Gibco, #A1443001) supplemented with 2% (w/v) fatty acid-free BSA (Sigma-Aldrich, #A8806)). On the day of the assay, the starvation medium was replaced with 50 µl fresh assay medium per adipoid. After 3 h, the medium was collected (basal lipolysis) and replaced with assay medium containing 10 µM isoproterenol (from a lipolysis assay kit; Akrivis Bio, #MA-0189). After another 3 h, the medium was collected to measure stimulated lipolysis. Glycerol was measured according to the manufacturer’s instructions using a standard curve of 0-4 nmol/well. Fluorescence was read at excitation/emission 535/587 nm on a FlexStation 3 microplate reader (Molecular Devices). Basal and stimulated glycerol release was calculated per hour per adipoid.

### Leptin and adiponectin immunoassays

Medium was collected from adipoids 72 h after a medium change, with the medium of three adipoids pooled per replicate, and stored at −20 °C until use. Adiponectin and leptin were measured with the Human Adiponectin ProQuantum Immunoassay Kit (Thermo Fisher Scientific, #A44838) and the Human Leptin ProQuantum Immunoassay Kit (Thermo Fisher Scientific, #A45250), respectively, according to the manufacturer’s instructions. Plates were read on a QuantStudio 6 (Thermo Fisher Scientific).

### Single-nucleus RNA sequencing

#### Nuclei isolation

Eight adipoids per line or donor were collected into 2 ml tubes (Eppendorf), washed twice in cold DPBS and snap-frozen in liquid nitrogen. Nuclei were isolated as described previously^21^. Briefly, frozen adipoids were transferred into a C-tube (Miltenyi, #130-093-237) containing 2 ml nuclei isolation buffer (Miltenyi, #130-128-024) supplemented with 1× cOmplete protease inhibitor (Sigma-Aldrich, #11697498001) and 0.5 units Protector RNase inhibitor (Sigma-Aldrich, #3335399001). The C-tube was placed in a gentleMACS dissociator (Miltenyi) at 4 °C, and nuclei were isolated using the 4C_nuclei program. The C-tubes were then incubated on ice for 5 min, and the nuclei suspension was filtered through a pre-wetted 30 µm filter (Sysmex, #04-004-2326) and mixed with 2 ml wash buffer (PBS, 1% (w/v) BSA, 1× cOmplete protease inhibitor, 0.5 U/µl RNase inhibitor). Nuclei were centrifuged at 500 × g and 4 °C for 5 min, and the pellet was resuspended in 1 ml wash buffer for counting.

### Library preparation

Nuclei were counted on a LUNA-FX7 automated cell counter (Logos Biosystems) using acridine orange/propidium iodide staining. Libraries were prepared using the GEM-X Universal 3′ v4 Gene Expression 4-plex assay (10x Genomics) according to the manufacturer’s instructions. We targeted 5,000 nuclei per sample per lane and ran all samples across two lanes, for a total of 10,000 targeted nuclei per sample. Libraries were sequenced on a NovaSeq X Plus (Illumina) with a read configuration of 28-10-10-90, targeting 35,000 reads per nucleus.

### Data analysis

Base calling was performed with bcl-convert (v4.3.13, Illumina), and FASTQ quality was assessed with FastQC (v0.12.1). Demultiplexing, read alignment to the GRCh38-2024A reference genome (10x Genomics), barcode processing and UMI counting were performed with Cell Ranger multi (v10.0.0, 10x Genomics). Datasets were processed, integrated and analyzed with the R package Seurat (v5.1.0)^22^. During quality control, nuclei were discarded if they had fewer than 300 or more than 6,000 detected features, more than 40,000 UMIs, or more than 5% mitochondrial or more than 5% ribosomal gene counts. The cluster resolution was determined for each dataset with the R package clustree^23^. Cluster markers were identified by differential gene expression analysis (Wilcoxon rank-sum test; |log2 fold change| > 0.25 and adjusted P < 0.01). Clusters with low feature and UMI counts but high mitochondrial and ribosomal gene counts were discarded. After clustering, nuclei were reference-mapped to the human white adipose tissue atlas^19^. Data from hiPSC-derived adipoids were first analyzed separately and then integrated with data from primary cell-derived adipoids using the RPCA method in Seurat, which was chosen after comparing CCA, RPCA and Harmony. For differential expression analysis between hiPSC-derived and primary cell-derived adipoids, pseudobulk count matrices were generated per cell type with the Seurat function AggregateExpression, restricted to genes expressed in at least 20% of nuclei in either condition, and analyzed with DESeq2 as implemented in Seurat. For correlation of pseudobulk gene expression, counts were log-normalized, and heatmaps were plotted with the R package ComplexHeatmap (v2.20.0). Visualizations were generated with Seurat v5, scCustomize (v3.2.4)^24^ and SCpubr (v3.0.1)^25^.

## Acknowledgements

We thank Kim Schneider and Michael Zaazman from the 360 Labs Genomics Platform for their support with generating snRNAseq libraries. We thank Mairene Coto Llerena, and Ece Yildiz for help in sourcing human adipose tissue samples.

## Author Contributions

L.C.H. and R.O. performed experiments. S.J. and L.C.H. generated snRNAseq libraries. S.E.S. S.T., and M.S. helped with cell culture experiments. M.S. and M.N.H. supervised the work. L.C.H., M.S., and M.N.H. wrote the manuscript. All authors contributed to reviewing and editing the manuscript.

---

**Table S1:** Patient data.

| Donor | BMI | Age | Sex |
|---|---|---|---|
| P13 | 26.7 | 28 | Female |
| P14 | 28 | 42 | Female |
| P15 | 28.9 | 32 | Male |

**Supplemental Figure S1.**
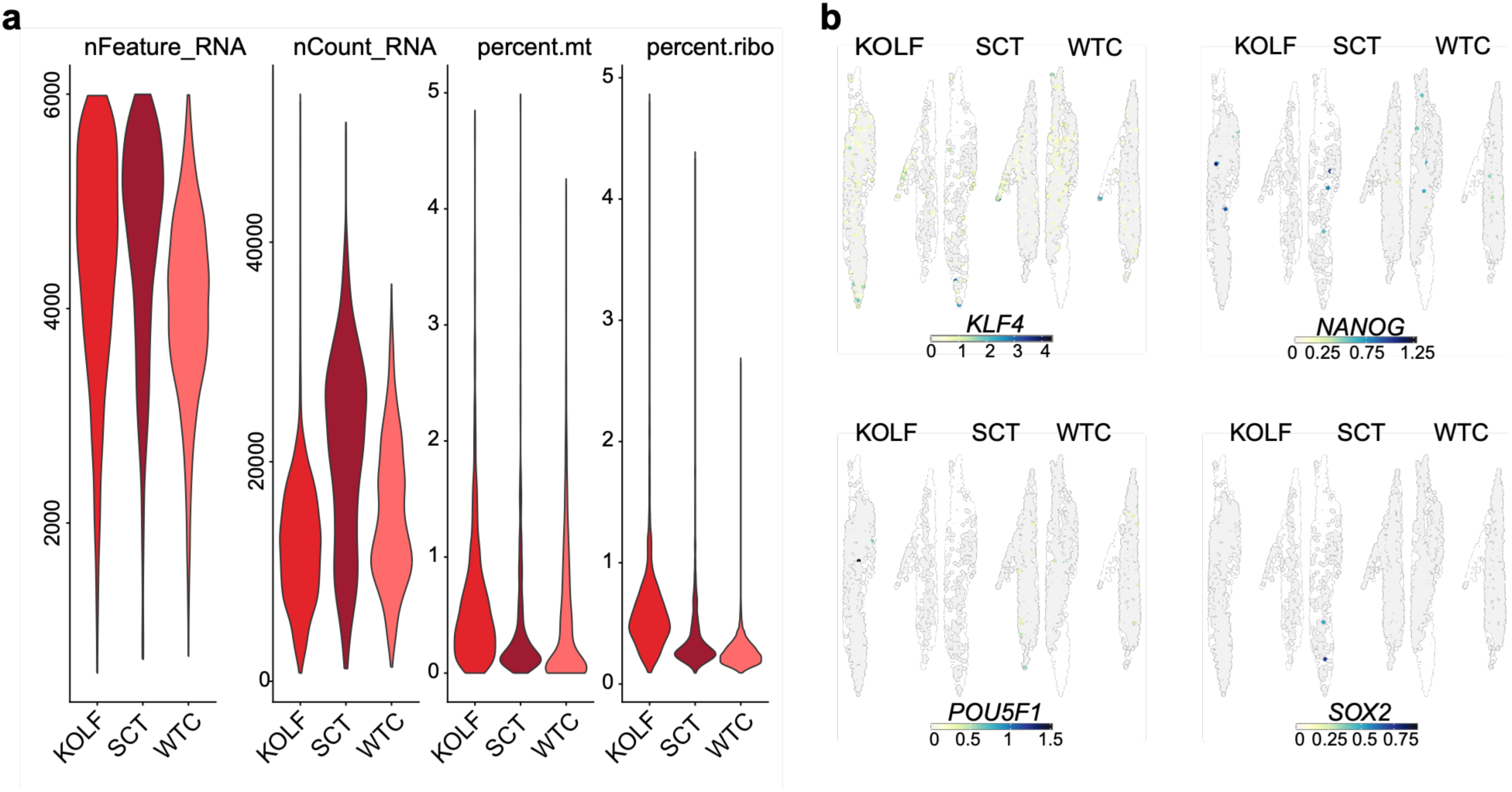
**a**, Gene counts, the number of UMIs, fraction of mitochondrial transcripts and fraction of ribosomal genes per nucleus per cell line. **b**, Normalized expression of pluripotency markers *KLF4*, *NANOG*, *POU5F1* and *SOX2* across adipoids from different hiPSC lines.

**Supplemental Figure S2.**
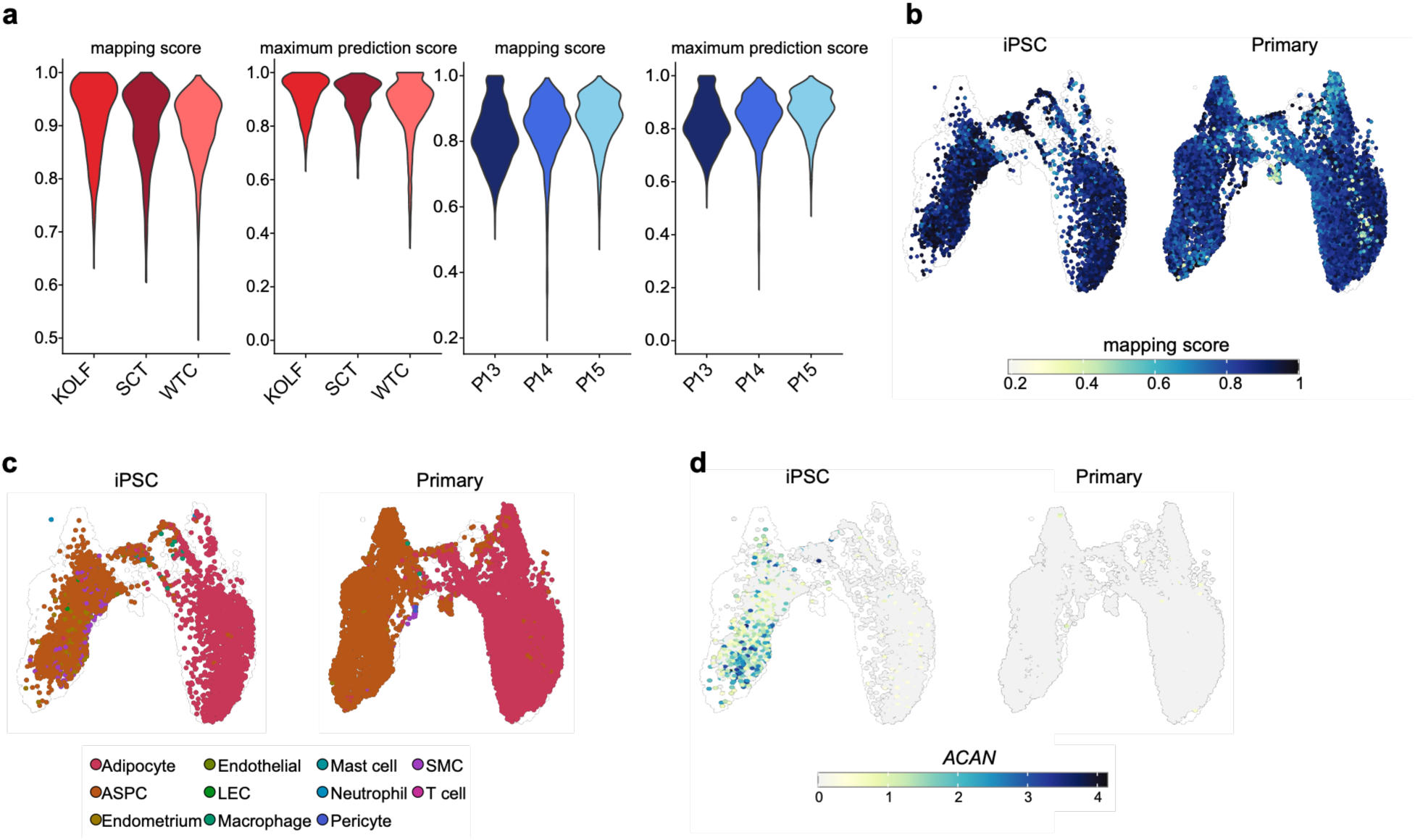
**a**, Mapping score and maximum prediction score distribution per cell line or donor adipoids obtained from reference mapping against Emont *et al.*^19^. **b**, Feature plot showing reference mapping scores of adipoid datasets projected onto to the Emont *et al.* ^19^. **c**, UMAP visualization representing hiPSC and primary cell derived adipoids colored by the predicted cell types from the Emont et al. human adipose tissue dataset. **d**, Normalized expression of *ACAN*, a marker gene for chondrogenic or myogenic lineages.

